# The breadth-depth dilemma in a finite capacity model of decision-making

**DOI:** 10.1101/2020.03.13.987081

**Authors:** Rubén Moreno-Bote, Jorge Ramírez-Ruiz, Jan Drugowitsch, Benjamin Y. Hayden

**Affiliations:** Center for Brain and Cognition, and Department of Information and Communication Technologies, Universitat Pompeu Fabra, Barcelona, Spain; Serra Húnter Fellow Programme, Universitat Pompeu Fabra, Barcelona, Spain; Department of Neurobiology, Harvard Medical School, Boston, MA 02115, USA; Department of Neuroscience, Center for Magnetic Resonance Research, and Center for Neural Engineering, University of Minnesota, Minneapolis, MN, 55455, USA

## Abstract

Decision-makers are often faced with limited information about the outcomes of their choices. Current formalizations of uncertain choice, such as the explore-exploit dilemma, do not apply well to decisions in which search capacity can be allocated to each option in variable amounts. Such choices confront decision-makers with the need to tradeoff between *breadth* - allocating a small amount of capacity to each of many options – and *depth* - focusing capacity on a few options. We formalize the breadth-depth dilemma through a finite sample capacity model. We find that, if capacity is smaller than 4-7 samples, it is optimal to draw one sample per alternative, favoring breadth. However, for larger capacities, a sharp transition is observed, and it becomes best to deeply sample a very small fraction of alternatives, that decreases with the square root of capacity. Thus, ignoring most options, even when capacity is large enough to shallowly sample all of them, reflects a signature of optimal behavior. Our results also provide a rich casuistic for metareasoning in multi-alternative decisions with bounded capacity.

## Introduction

The breadth-depth (BD) dilemma is a ubiquitous problem in decision-making. Consider the example of going to graduate school, where one can enroll in many courses in many topics. Let us assume that the goal is to determine the single one topic that is most relevant, the one that will grant us a job. Should I enroll in few courses in many topics –breadth search— at the risk of not learning enough about any topic to tell which one is the best? Or should I enroll in many courses in very few topics –depth search— at the risk of missing the really exciting topic for the future? One crucial element of this type of decision is that the allocation of resources (time, in this case) needs to be done in advance, before feedback is received (before classes start). Also, once decided, the strategy cannot be changed on the fly, as doing so would be very costly. The BD dilemma is popular in tree search algorithms (Horowitz and Sahni, 1978; Korf, 1985) and in optimizing menu designs (Miller, 1981). It is also one faced by humans and other foragers in many situations, as when we plan, schedule, or invest with finite resources. It is remarkable that the bulk of research on the BD has been in fields outside of psychology (e.g. (Halpert, 1958; Schwartz et al., 2009; Turner et al., 2002). We believe that one reason is the lack of standard sets of tools for thinking about the problem and separating it from other dilemmas.

Many features of the BD dilemma warrant its study in isolation. First, BD decisions are about how to divide finite resources, with the possibility of oversampling specific options and ignoring others. Second, the BD dilemma is about making strategic decisions, that is, decisions that need to be planned in advance and cannot be changed on the fly once initiated, e.g., once courses start, it is very costly to change them. Finally, these decisions need to be made before feedback is received: enrollment happens before courses start, and thus before knowing the true relevance of the courses and topics. These features are distinct from those of the well-known exploitation-exploration dilemma (Cohen et al., 2007; Costa et al., 2019; Daw et al., 2006; Ebitz et al., 2018; Wilson et al., 2014) and its associated formalization in multi-armed bandits (Averbeck, 2015; Chen et al., 2016; Gittins et al., 2011).

Past work revealed that humans appear to carefully trade off the benefits of breadth and depth in multi-alternative decision-making. For example, when faced with a large number of options, we often focus – even if arbitrarily – on a subset of them (Bettman et al., 1998; Brandstätter et al., 2006; Gigerenzer and Gaissmaier, 2011; Tversky, 1972) with the presumable benefit that we can more precisely evaluate them. Likewise, we may consider all options, but arbitrarily reject value-relevant dimensions (Busemeyer et al., 2019; Timmermans, 1993), as if contemplating them all is too costly. Option narrowing appears to be a very general pattern, one that is shared with both human and non-human animals, despite the fact that rejecting options can reduce experienced utility (Gigerenzer and Gaissmaier, 2011; Tversky, 1972). It is often proposed that such heuristics reflect bounded rationality (Simon, 1955), which is likely correct in principle, but the exact processes underlying that rationality remain to be identified. Why do we so often consider a very small number of options when considering more would a priori improve our choice? One possibility is that this pattern reflects an evolved response to an empirical fact: that when capacity is constrained, optimal search favors consideration of a small number of options.

Because cognitive capacity is limited in many ways, the BD dilemma has direct relevance to many aspects of cognition as well. For example, executive control is thought to be limited in capacity, such that control needs to be allocated strategically (Hills et al., 2010; Koechlin and Summerfield, 2007; Shenhav et al., 2013, 2017). Likewise, attentional focus and working memory capacity are limited, such that, during search, we often foveate only a single target or hold a few items in memory (Cowan et al., 2005). Although the effective numbers are low, each contemplated option is encoded with great detail (Awh et al., 2007; Luck and Vogel, 2013; Ma et al., 2014). Furthermore, it seems clear that recollection of information from memory can be thought of as a search-like process (Hills et al., 2012; Ratcliff and Murdock, 1976; Shadlen and Shohamy, 2016). That is, to retrieve a memory we must attend to a recollection processes, with its associated limited capacity. Thus memory-guided decisions presumably involve BD tradeoffs too.

Although the relevance of the BD dilemma is clear, tractable models are lacking, and thus, optimal strategies for BD decisions are largely unknown. Here, we develop and solve a model for multi-alternative decision making endowed with the prototypical ingredients of the BD dilemma. Our model consists of a reward-optimizing yet bounded decision-maker (Gershman et al., 2015; Griffiths et al., 2015; Simon, 1955) confronted with multiple alternatives with unknown subjective values. The first critical element of the model is *finite sample capacity*, which enforces a tradeoff between sampling many options with few samples (breadth) and sampling few options with many samples (depth). The second critical element is that samples need to be allocated across alternatives before sampling starts and, thus, before feedback is available. This strategic decision with the finite sample capacity constraint implies a metareasoning problem (Griffiths et al., 2015; Russell and Wefald, 1991) where deliberation about the multiple possible allocations of resources (meta-actions) need to be made in advance to optimize expected utility of a future choice.

Despite the simplicity of the model, it features non-trivial behaviors, which are characterized analytically. When capacity is low (less than 4-7 samples can be probed), it is best to sample as many alternatives as possible, but only once each; that is, breadth search is favored. At larger capacities, there is a qualitative and sharp change of behavior (a ‘phase transition’) and the optimal number of sampled alternatives grows with the square root of sample capacity (‘square root sampling law’), balancing breadth and depth. Therefore, in this regime it is best to ignore the vast majority of potentially accessible options. We considered globally optimal allocations in comparison to even allocation of samples across sampled alternatives and found that the square root sampling law, obtained for the latter, provides a close-to-optimal heuristic that is simpler to implement. Our results are robust to strong variations of the environments where the probability of finding good options widely varies.

## Results

### Finite sample capacity model

We assume that a decision-maker can choose how to allocate a finite resource among options of unknown status to determine the best option (**Figure 1**). The environment generates a large number of options, each characterized by the probability of delivering a successful outcome. The success probabilities, unknown to the decision-maker, determine the quality of each of the options, with better options having higher success probabilities (e.g., options with a higher probability of delivering a large reward if they are sampled). The goal of the decision-maker is to infer which of the options has the highest success probability. The success probabilities of the options are generated randomly from an underlying prior probability distribution, modelled as a beta distribution. We assume that this distribution is known by the decision-maker due, for example, to previous experience with the environment. The prior distribution defines the overall difficult of finding successful options in the environment.

**Figure 1.**
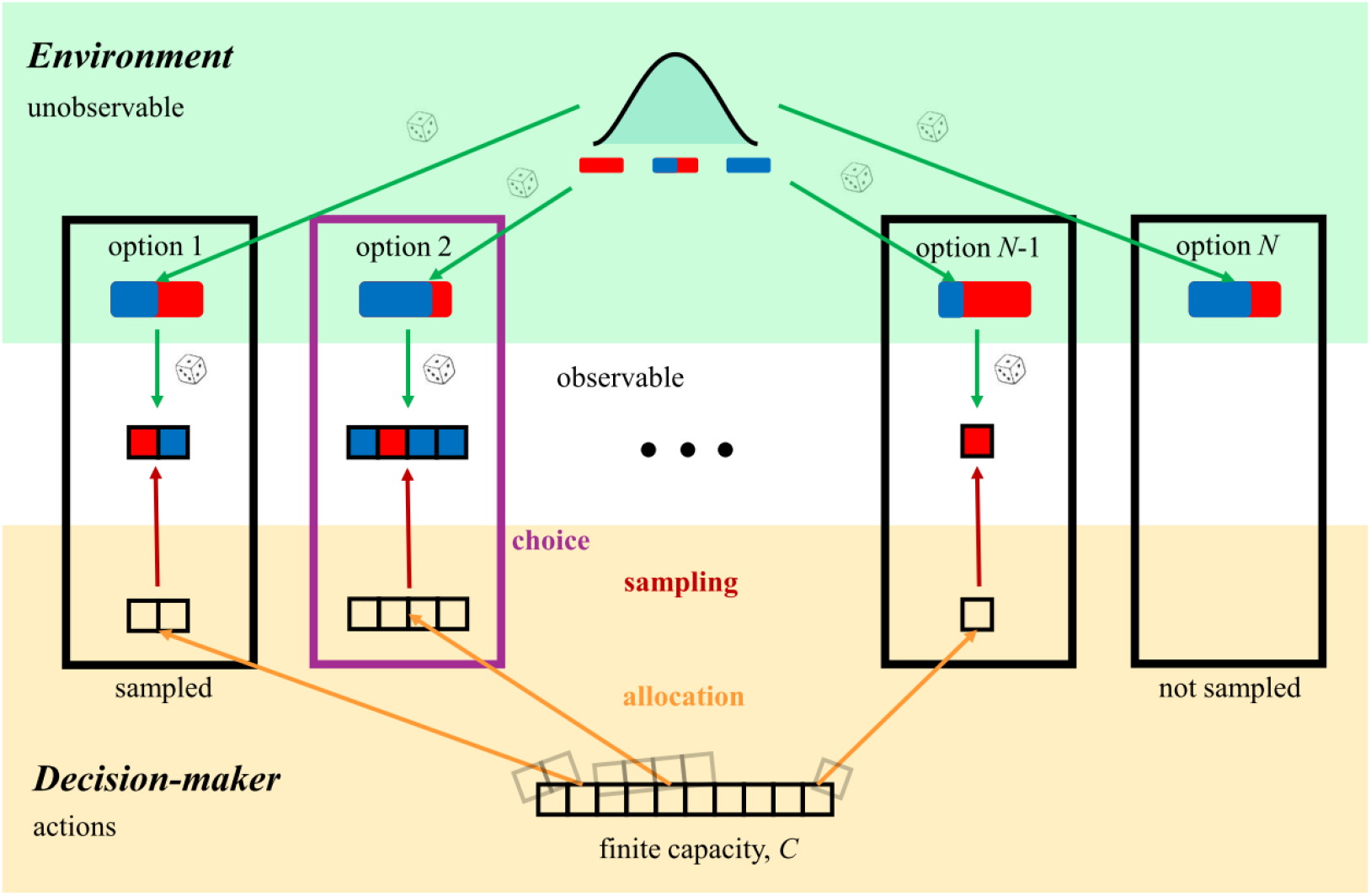
Finite sample capacity model. The environment (top, green) contains a large number *N* of options, and choosing either might lead to a successful outcome (e.g., a large reward). For each option, the probability of success (blue fraction of red/blue bar) is a-priori unknown to the decision-maker, and is drawn independently across options from an underlying prior probability distribution, modelled as a beta distribution (top distribution). The prior distribution defines the overall difficult of finding successful options in the environment. Options are characterized by the probability of delivering a successful outcome (e.g., a large reward), and the outcomes are modelled as Bernoulli variables. The decision-maker (bottom, orange) has a finite capacity *C*, i.e., a finite number of samples (bar of squares) that can be allocated to any option in any possible way. The decision-maker can decide to oversample options by allocating more than one sample to them (e.g., options on the left), and also ignore some options by not sampling them at all (e.g., rightmost option). All samples need to be allocated in advance and allocation cannot be changed thereafter. Therefore, feedback is not provided at this stage. After allocation, sampling starts, in which the decision-maker observes a number of successes and failures for each of the sampled options (colored squares; blue: success –large reward, red: failure –small reward). Once this evidence is collected, the decision-maker choses the option that is deemed to have the highest probability of success (in this case, option 2; purple box).

The decision-maker is endowed with a finite sample capacity *C*, i.e., a finite number of samples that she can allocate to any option and to as many options as desired. Within the allowed flexibility, it is possible that the decision-maker decides to oversample options by allocating more than one sample to them, and it is also possible that she decides to ignore some options by not sampling them at all. Feedback is not provided at the allocation stage, so this decision is based purely on the expected quality of options in the environment. After allocation has been determined, the outcomes of the samples are revealed, constituting the only feedback that the decision-maker receives about the fitness of her sample allocation. Outcomes for each of the sampled alternatives are modelled as a Bernoulli variable, where a successful outcome (corresponding to a large reward) has probability equal to the success probability of that option. The inferred best alternative is the one with the largest inferred success probability based on the observed outcomes from the allocated samples to each of the options (Bechhofer and Kulkarni, 1984; Gupta and Liang, 1989; Sobel and Huyett, 1957). Choosing this alternative maximizes expected utility (see below and Methods).

While making a choice based on the observed outcomes is a trivial problem, deciding how to allocate samples over the options to maximize expected future reward is a hard combinatorial problem. There are many ways a finite number of samples can be allocated amongst a very large number of alternatives. At the *breadth* extreme, one can split capacity to over as many alternatives as possible, sampling each just once. In this case, the decision-maker will likely identify a few promising options, but will lack the information for choosing between them. At the *depth* extreme, the search could allocate all samples to one alternative. The decision-maker’s estimate of the success probability of that option will be accurate, but that of the other alternatives will remain unknown. It would seem that an intermediate strategy is better than either extreme. Specifically, the optimal allocation of samples should balance the diminishing marginal gains of sampling a new alternative and those of drawing an additional sample from an already sampled alternative.

To formalize the above model, let us assume that the decision-maker can sample and choose from *N* = *C* alternatives. That is, we consider scenarios where the number of alternatives *N* is as large as the decision-maker’s sampling capacity –if the number of alternatives is larger than capacity, the only difference is that there would be a larger number of ignored alternatives. The allocation of samples over the alternatives is described by the vector 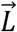, with components *L*_*i*_ representing the number of samples allocated to alternative *i* = 1, …, *N*. The finite sample capacity of the decision-maker imposes the constraint ∑_*i*_ *L*_*i*_ = *C*. Denoting *n*_*i*_ as the number of successes (1’s) of the Bernoulli variable over the *L*_*i*_ samples drawn from alternative *i*, the decision-maker’s expected utility 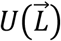 is

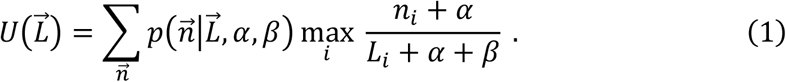

The optimal allocation of samples across options 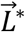 is the one that maximizes the decision-maker’s expected utility 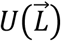 over all allocations of samples 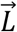,

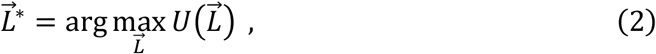

with the above finite sample capacity constraint (see Methods for details). The right-hand side in Eq. (1) results from taking the average over the expected gain associated with each possible number of successes 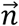, weighted by the probability of these successes occurring for a chosen allocation of samples 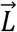. As each alternative is sampled independently, the joint distribution of success counts factorizes as 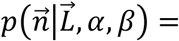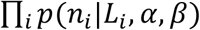, where *p*(*n*_*i*_|*L*_*i*_, *α*, *β*) is a beta-binomial distribution (Murphy, 2012), representing the probability of having exactly *n*_*i*_ successes from a Bernoulli variable that is drawn *L*_*i*_ times, and whose success probability *p*_*i*_ follows a beta distribution with parameters *α* and *β*. These parameters control the skewness of the distribution: with equal values of both parameters, the distribution is symmetric around one half, while for *α* larger (smaller) than *β* the distribution is negatively (positively) skewed. Overall, these parameters describe the difficulty of finding successful options in the environment. The term 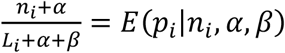 in Eq. (1) corresponds to the posterior mean for the beta-binomial distribution, and thus it represents the expected success probability of the sampled Bernoulli variable *i* based on the observed outcome and the prior distribution. The optimal expected utility then becomes 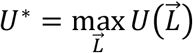, which involves a double maximization over the expected success probabilities of the sampled alternatives and the allocation of samples over the alternatives, effectively solving the two-stage decision process (i.e., first allocate samples, then observed outcomes, then choose) in reverse order (i.e., first optimize choices given outcomes and allocation, then optimize allocation).

This maximization allows total flexibility over how many samples to allocate to each alternative. For tractability, let us first consider the best even allocation of samples, that is, a subfamily of allocation strategies where the same number of samples *L* allocated to each of *M* sampled alternatives, while the remaining alternatives (*C* − *M*) are not sampled, subject to the standard capacity constraint *M* × *L* = *C*. Indeed, finding the optimal even allocation of samples is easier than finding the globally optimal allocation, which might be uneven in general (see below). As we show in Methods, a particularly simple expression for the optimal even sample allocation, *L*\*, arises when the prior distribution over success probabilities is uniform (*α* = *β* = 1),

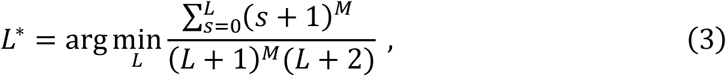

where the right-hand side is related to utility by

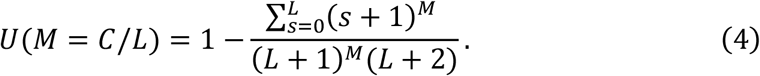

Note that only *M*\* = *C*/*L*\* ≤ *C* alternatives are sampled in the optimal allocation, while the remaining options are given zero samples, thus effectively being ignored. The sampled alternatives can be chosen randomly, as they are indistinguishable before sampling. By using extreme value theory (see Methods) we show that the optimal number of sampled alternatives *M*\* and optimal number of samples per alternative *L*\* both follow a power law with exponent ½ for large capacity *C*

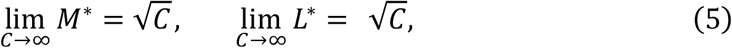

which corresponds to perfect balancing breadth and depth.

In the next section, we analyze this case in detail. After that, we consider optimal even allocations of samples for arbitrary prior distributions, and finally we provide results for the globally optimal allocations, not necessarily even.

### Sharp transition of optimal sampling strategy at capacity equal to 7 samples

We first analyze the expected utility *U*(*M*) as a function of the number of sampled alternatives *M* evenly each *L* times (such that *M* × *L* = *C*) (**Figure 2a**). At low capacity (*C* = 4, lighter gray line), the utility increases monotonically from sampling just one alternative (*M* = 1) four times, to sampling four alternatives (*M* = 4) one time each. Thus, a pure breadth strategy is favored. At intermediate capacity (*C* = 10, medium gray line), the maximum occurs at an intermediate number of alternatives (specifically, *M*=5), reflecting an increasing emphasis on depth. At large capacity (*C* = 100, black line), the maximum expected utility occurs when sampling few different alternatives (*M* = 10 sampled alternatives with *L* = 10 samples each), reflecting a tight balance between breadth and search. For such large capacities, sampling instead most of the of alternatives (rightmost point of the black line) would lead to a reward that approaches 2/3, which is the lowest expected reward one would obtain if at least one sampled alternative has a positive outcome (see Methods).

**Figure 2.**
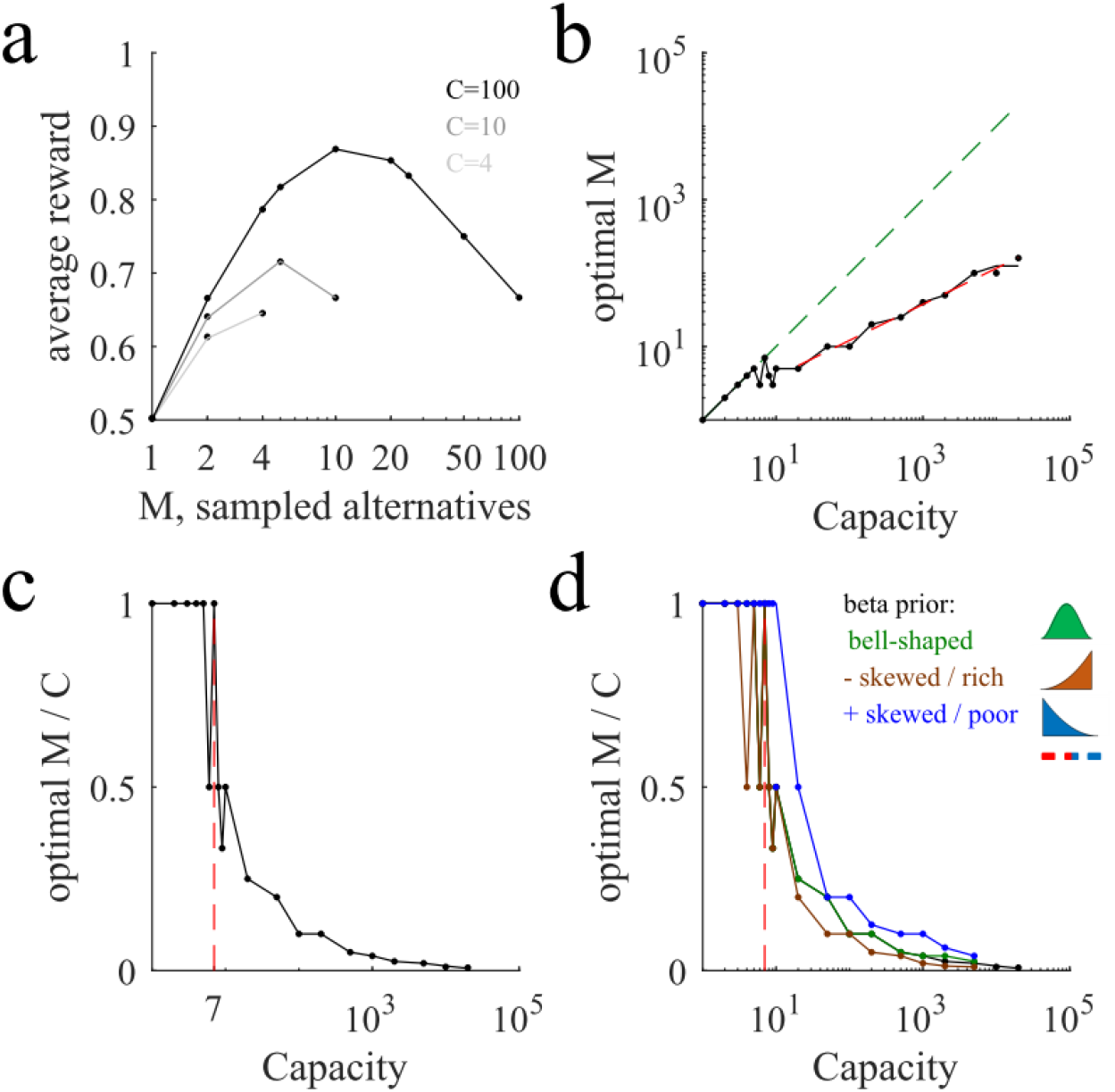
Sharp transitions in optimal number of sampled alternatives when crossing around the critical capacity of 4-7. (**a**) Average reward (points, simulations; lines, theoretical expressions, Eq. 4) as a function of the number of sampled alternatives *M* for three different capacities (*C* = 4, 10, 100; light, intermediate and dark lines respectively). The maximum occurs at the large extreme for low capacity but at a relatively low value for large capacity. Note log horizontal scale. (**b**) Optimal number of sampled alternatives as a function of capacity. When capacity is smaller than around 7, a linear trend of unit slope is observed (dashed green line), but when capacity is above 7, a sublinear behavior is observed (dashed red line corresponds to the best power law fit, with exponent close to ½). The transition between these two regimes is sharp. Black line corresponds to analytical predictions –the jagged nature of this prediction and simulation lines in this and other panels is due to the discrete values that the optimal *M* can only take, not to numerical undersampling. (**c**) The sharp transition is clearer when plotting the optimal number of sampled alternatives to capacity ratio as a function of capacity. For low capacity, the ratio is one, but for large capacity the ratio decreases very rapidly. The last point for which the optimal ratio is one corresponds to capacity equal to 7 (indicated with a vertical red line). (**d**) Number of sampled alternatives to capacity ratios for different prior distributions (*α* = *β* = 1, black line, same as in the previous panel, not visible because it is very similar to the one with a bell-shaped prior; *α* = *β* = 3, bell-shaped, green line; *α* = 3, *β* = 1, negatively skewed prior modelling a ‘rich’ environment, brown line; *α* = 1, *β* = 3, positively skewed distribution modeling looking for a ‘needle in a haystack’, that is, a ‘poor’ environment, blue line). Lines correspond to analytical predictions from Eq. 3; points correspond to numerical simulations; error bars are smaller than data points in all panels.

The model displays a sharp transition when capacity crosses the critical value of 7 (**Figure 2b**). Below this transition point, the optimal number of sampled alternatives equals capacity –except for capacity 6, where there is a temporary dip below one. That is, one should follow a breadth strategy and distribute one sample to each alternative. Above 7, the optimal number of sampled alternatives is much smaller than the capacity. That is, one should balance the number of sampled alternatives with the depth of sampling each of them. Specifically, the optimal number of sampled alternatives follows a power law with exponent ½ (log-log linear regression, power = slope = 0.50, 95% CI = [0.48, 0.52]), as predicted by Eq. (5), which implies that the fraction of sampled alternatives decreases with the square root of capacity. This means that breadth and depth are tightly balanced in the optimal strategy. The sharp transition at 7 becomes clearer when plotting the ratio between the optimal number of sampled alternatives and capacity as a function of capacity (**Figure 2c**).

In summary, if the capacity of a decision-maker increases by a factor of 100, the decision-maker will roughly increase the number of samples alternatives just by a factor of 10, one order of magnitude smaller than the capacity increase. Because the optimal number of sampled alternatives increases with capacity with an exponent ½, we call this the ‘square root sampling law’. A remarkable implication of this law is that the vast majority of potentially accessible alternatives should be ignored (e.g., for *C* = 100, *C* − *M* = 90 options are ‘rationally’ ignored).

### Generalizing to variations in beta prior distributions

The above critical capacity for optimal even sample allocation changes when, instead of using a uniform prior of success probabilities, we allow for variations of the prior distribution (e.g. **Figure 2d**), but it consistently lies in the range 4-7 with the critical value depending on the environment. By changing the prior’s parameters, we can vary the difficulty of finding a good extreme alternative, and thus can compare different scenarios. For the uniform prior that we have used above, a decision-maker is equally likely to find an alternative with any success probability. Consider a prior distribution that is concentrated and symmetric around a success probability of 0.5 (approximately as a Gaussian, corresponding to the beta prior parameters *α* = *β* = 3). In this environment, unusually good (high success probability) and unusually bad (low success probability) options are rarer than medium ones (**Figure 2d**, green line). In this case, the breadth-depth tradeoff as a function of *C* is remarkably similar to the uniform prior case, with a transition at *C* = 7.

We also consider a negatively skewed prior distribution (*α* = 3, *β* = 1). This distribution refers to environments with rare bad options, as, for example, a tree whose fruits are mostly ripe but that has a few unripe ones. In this ‘rich’ environment, one can afford sampling a smaller number of options, but as they are sampled more deeply, it is possible to detect better the really excellent ones. A sharp transition occurs even in this condition, and the last point for which the ratio is one corresponds again to the critical capacity of 7 (brown line), while capacity 4 corresponds to the first point where the ratio is below one. As expected in this environment, the decay of the ratio after this transition is (slightly) faster than that of the symmetric prior. Therefore, negative skews engender a modest bias towards depth over breadth.

Finally, consider the opposite scenario, in which the prior distribution is concentrated at low success probability values (*α* = 1, *β* = 3, positively skewed beta distribution), which corresponds to looking for a ‘needle in a haystack’ or a ‘poor’ environment. In this scenario, one ought to sample more alternatives less deeply to allow for the possibility of finding the rare good alternatives, and thus breadth should be emphasized over depth (**Figure 2d**, blue line). In this scenario, the sharp transition occurs at capacities around 10 (blue line).

Despite the large variations of prior distributions, a fast transition occurs in all conditions at around a small capacity value, as in the uniform prior case. In addition, a power law behavior is observed for values of *C* greater than around 7 regardless of skew, with exponents close to ½ in all cases (uniform prior, power = 0.50; negatively skewed prior, 0.46; positively skewed prior, 0.63; s.e.m. < 0.02).

### Optimal choice sets and sample allocations

So far, we have studied optimal even sample allocation. Let us now consider the payoffs for decision-makers willing to consider all possible allocation strategies. The number of all possible allocations equals the number of partitions of integers in number theory, which grows exponentially with the square root of capacity (Andrews, 1998). This makes finding the globally optimal sample allocation a problem that is intractable in general. For small capacity values *C* ≤ 7 and uniform prior distributions we compute the exact optimal sample allocation by exhaustive search and rely on a Monte Carlo gradient descent method for larger capacities and other priors. The latter finds a local maximum for the reward, that we found to coincide with the exhaustive search optima for small capacities.

Globally optimal sample allocation (which defines optimal choice sets) for a uniform prior beta distribution tends to sample all or most of the alternatives when the capacity is small, but as capacity increases the number of sampled alternatives decrease (**Figure 3**, left). For instance, for capacity equal to 5 samples, the optimal sample allocation is (2,1,1,1,0). In general, in optimal allocations, the decision-maker adopts a local balance between oversampling a few alternatives and sparsely sampling others –a local compromise between breadth and depth— even though all options are initially indistinguishable.

**Figure 3.**
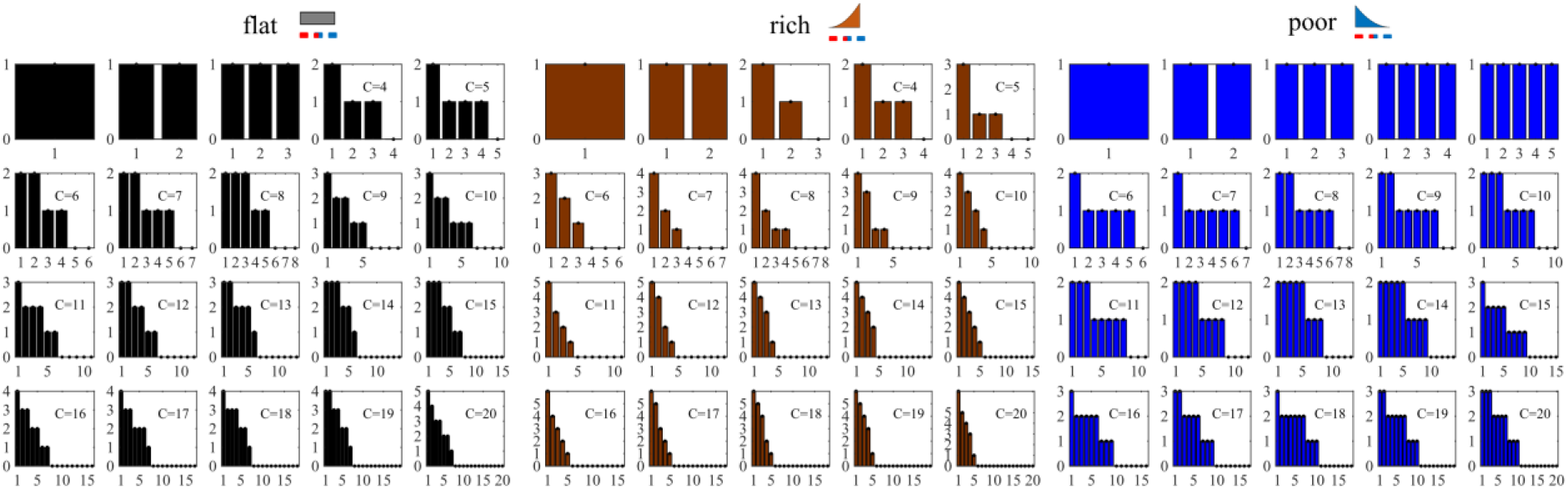
Optimal sample allocations and choice sets. Optimal sample allocation for flat, rich and poor environments from capacity *C* = 1 up to *C* = 20. The environments corresponding respectively to uniform, negatively and positively skewed prior distributions (top icons). Optimal sample allocations are represented as bar plots, indicating the number of samples allocated to each alternative ordered from the most to the least sampled alternative. Note that at large capacity many alternatives that could potentially be sampled (up to a number equal to the capacity) are not actually sampled.

We also studied optimal sample allocation for positively and negatively skewed prior distributions. In a rich environment (center panel), the optimal sample allocation is uneven for capacities as low as *C* = 3. In contrast, in a poor environment (right), the optimal sample allocation remains even up to capacity *C* =5, which was not the case for the flat environment (compare with left panel). For higher capacities, decision-makers in rich environments ought to sample less broadly but more deeply. For instance, for capacity *C* = 20, only around 5 alternatives are sampled, while the remaining 15 potentially accessible alternatives are neglected. In the haystack environment, in contrast, about half of the alternatives are sampled, but not very deeply (only a maximum of 3 samples are allocated to the most sampled alternatives).

### Even sample allocation is close to optimal

Three principles stand out. First, globally optimal sample allocation almost never coincides with optimal even allocation. Second, at low capacity optimal allocation favors breadth while at large capacity a breadth-depth balance is preferred (**Figure 4a**). Third, a fast transition is observed between the two regimes happening at a relatively small capacity value. The last two features are shared by the optimal even allocations as well (cf. Figure 2c).

**Figure 4.**
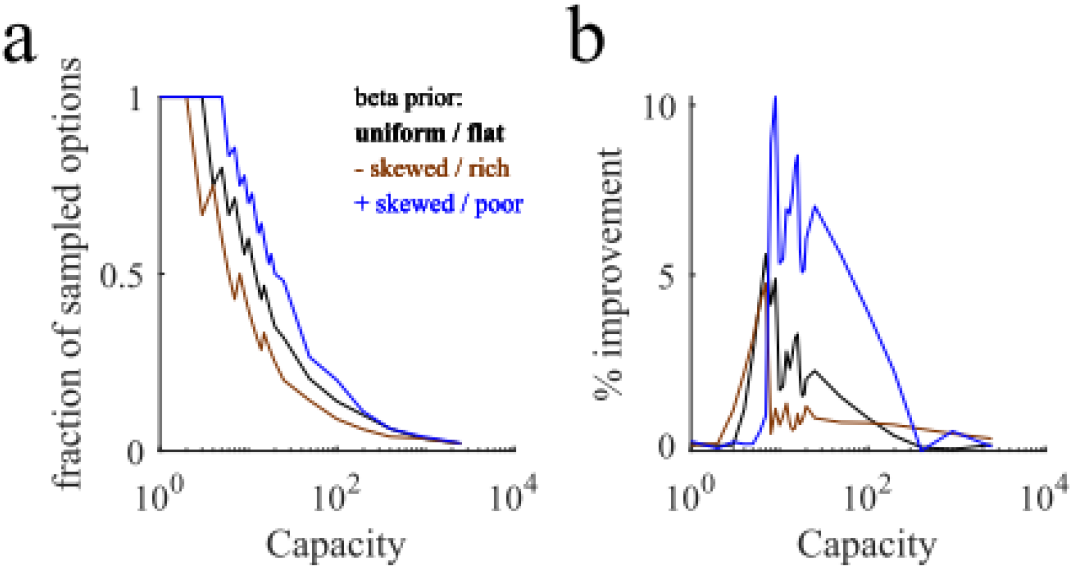
Globally optimal and optimal even sample allocations share similar features and have similar performances. (**a**) Fraction of sampled options (compared to the maximum number of potentially accessible alternatives, equal to *C*) as a function of capacity *C*. The fraction is close to one for small values for all environments (flat -black line, rich -brown, and poor -blue). The fraction decays rapidly to zero from a critical value that depends on the prior. The jagged nature of the lines is due to the discrete nature of capacity. (**b**) Percentage points increase in averaged reward by using globally optimal sample allocation compared to even allocation (see Methods). Color code as in the previous panel.

Optimal even and globally optimal sample allocations share some important features, but are they equally good in terms of average reward obtained? We compared the average reward from globally optimal and even optimal sample allocations. For comparison, we always used optimal even sampling based on a uniform prior over each alternatives’ success probabilities, that is, we sample *M* = *C* alternatives with one sample each if capacity is *C* ≤ 7 and 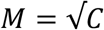 alternatives with 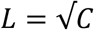 samples each if capacity is larger (square root sampling law; see Methods for details). This heuristic produced comparable performances to the optimal ones (**Figure 4b**). The worst-case scenario occurred in the poor environment (blue line) when capacity is close to 10, which led to a reduction in performance of close to 10%, but the maximum discrepancy value was even smaller for the flat and rich environments. Indeed, for the flat environment, the maximum reduction of performance was only around 5%.

For large capacity *C* > 100 the square root sampling law produced results that were very close to the performance of the optimal solutions (as found by the MC gradient descent method). Therefore, the advantage provided by using globally optimal sample allocation over optimal even sampling at low capacity and the square root sampling law for high capacity is at most marginal.

## Discussion

We delineate a formal mathematical framework for thinking about a commonplace decision-making problem. The *breadth-depth dilemma* occurs when a decision-maker is faced with a large set of possible options, can query multiple options simultaneously, and has a limited search capacity. In such situations, the decision-maker will often have to balance between allocating search capacity to more (breadth) or to fewer (depth) options. We develop and use a finite sample capacity model to analyze optimal allocation of samples as a function of capacity. The model displays a sharp transition of behavior at a critical capacity close to the magical numbers 4-7 (cf. (Miller, 1956)). Below this capacity, the optimal strategy is to allocate one sample per alternative to access to as many alternatives as possible (i.e. breadth is favored). Above this capacity, breadth-depth balance is emphasized, and the *square root sampling law*, a close-to-optimal heuristic, applies. That is, capacity should be split into a number of alternatives equal to the square root of the capacity. This heuristic provides average rewards that are close to those from the optimal allocation of samples. As it is easy to implement, it can become a general rule of thumb for strategic allocation of resources in multi-alternative choices. The same results roughly apply to a wide variety of environments, including flat, rich and poor ones, characterized by very different difficulties of finding good options.

Despite the billions of neurons in the brain, our processing capacity seems quite limited. This strict limit applies to attention (where it is sometimes called the attentional bottleneck, (Deutsch and Deutsch, 1963; Treisman, 1969; Yantis and Johnston, 1990), including spatial attention, where the limit is best characterized (Desimone and Duncan, 1995), over working memory (Brady et al., 2011; Cowan et al., 2005; Luck and Vogel, 2013; Ma et al., 2014; Miller, 1956; Sims, 2016), to executive control (Norman and Shallice, 1986; Shenhav et al., 2017; Sleezer et al., 2016) and to motor preparation (Cisek and Kalaska, 2010). These narrow limits, which often number only a handful of items (though see (Ma et al., 2014)), suggest some sort of bottleneck. However, another interpretation is that capacity is much larger than it appears, and instead, observed capacity reflects the strategic allocation of resources according to the compromises that our model identifies as optimal. The square root sampling law, in other words, suggests that the apparently narrow bandwidth of cognition may reflect the optimal allocation across very few options of a relatively large capacity.

This is particularly likely to be true for economic choice. We are especially interested in the apparent strict capacity limits of prospective evaluation (Hayden and Moreno-Bote, 2018; Krajbich et al., 2010; Lim et al., 2011; Redish, 2016; Rich and Wallis, 2016). Indeed, the failures of choice with choices sets over a few items are striking and have been a major part of the heuristic literature (Diehl and Poynor, 2010; Iyengar and Lepper, 2000). These strict limits are ostensibly difficult to explain. It does not derive, for example, from the basic computational or biophysical properties of the nervous system, as is evident from the fact that our visual systems are an exception to the general pattern and can process much information in parallel. Nor do these limits appear to relate to any desire to reduce the extent of computation, as large numbers of brain regions coordinate to implement these cognitive processes (Rushworth et al., 2011; Siegel et al., 2015; Vickery et al., 2011; Yoo and Hayden, 2018). Our results presented above offer an appealing explanation for this problem: economic choice can be construed as breadth-depth search problems, and even when capacity is large, the optimal strategy is to focus on a very small region of the search space. Thus, our results can also help to understand why many cognitive systems operate in a regime of low sampling size, thus resolving the paradox of why low breadth sampling and large brain resources can coexist. We believe that these results are particularly relevant to behavioral economics.

Research has shown that consumers often consider just a small number of brands from where to purchase a specific product out of the many brands that exist in the market (Hauser and Wernerfelt, 1990; Stigler, 1961). The prevailing notion is that decision-makers hold a consideration choice set from where to make a final choice rather than contemplating all possibilities. Several reasons for this behavior have been provided. First, choice overload has been shown to produce suboptimal choices in certain conditions (Iyengar and Lepper, 2000; Scheibehenne et al., 2010). Secondly, selecting a small number of options from where to choose can be actually optimal if there is uncertainty about the value of the options and there is cost for exploring and sampling further options (Mehta et al., 2003; Roberts and Lattin, 1991; Santos et al., 2012).

Estimating the overall benefits of considering larger sets has to be balanced with the associated cost of exploring further options. This research has provided a relevant line of thought to understand low sampling behavior within the context of bounded rationality by formally assuming the presence of linear costs of time for searching for new options. Time costs comes in the model at the expense of unknown parameters, which often are difficult to fit (Mehta et al., 2003; Roberts and Lattin, 1991). Further, linear time costs always permit unlimited number of sampled options, as they do not impose a strict limit in the number of options that can be sampled. In our approach, in contrast, allocating finite resources imposes a strict limit to the number of options that can be sampled and, as resources are limited, there is a tradeoff between sampling more options with less resources or sampling fewer options with more resources, directly addressing the breadth-depth dilemma. This difference could be the main reason why the consideration set literature has not reported sharp transitions of behavior as a function of model parameters (costs) nor power sampling laws, which are the main features of our finite sample capacity model.

A number of extensions would be required to fully address more realistic problems associated to the breadth-depth dilemma. So far, we have considered a two-stages decision process, where the first metareasoning decision is about optimally distributing limited sampling capacity. It would be interesting to extend our results to sequential processes, where the decision of how to allocate is iterated over several steps, with intermediate feedback. An advantage of this more general setup (Morgan and Manning, 1985) is that a full-fledged interaction between the breadth-depth and exploration-exploitation dilemmas could be studied. In particular, a relevant direction is relating our square root sampling law with Hick’s law (Hick, 1952) for multialternative choices. The two approaches touch different aspects of multialternative decision making: while Hick’s law refers to the problem of how long options should be sampled in a multialternative setting, it does so by sampling all available options; the square root sampling law, by contrast, applies to situations where there are many alternatives and large fraction of them are to be ignored due to limited capacity, directly facing the breadth-depth dilemma. It will be interesting to integrate the two sets of results within a general framework of multialternative sequential sampling (Roe et al., 2001; Tajima et al., 2016; Usher and McClelland, 2004) under limited resources.

A second possible extension of our work is reconsidering the nature of capacity. For instance, ‘rate distortion theory’ defines a natural capacity constraint over the mutual information between the inputs and the outputs in a system (Bates et al., 2019; Sims, 2003, 2016). This capacity constraint might more naturally enforce a finite capacity than fixing the total number of samples that a system can draw from (externally or internally). A third relevant direction would be extending our study to cases where the capacity is continuous rather than discrete, and to cases where the observations are continuous variables. Although this remains a topic for future research, we do not expect qualitative differences in behavior, as for large capacity the continuous limit approximation applies, and for low capacities a low number of sampled alternatives is expected.

While we do not know of direct tests of breadth-depth capacities in humans, indirect measurements suggest that the square root sampling law can be at work in some realistic conditions, such as chess. It has been argued that chess players can image around 100 moves before deciding their next move (Simon, HA, 1972). Assuming that their capacity is 100, then the square root sampling law would predict that players should sample 10 immediate moves followed by around 10 continuations. Indeed, estimates indicate that chess players mentally contemplate roughly between 6-12 immediate moves followed by their continuations (Simon, HA, 1972) before capacity is exhausted due to time pressure. Although decisions in trees like this surely involve other types search heuristics beyond balancing breadth and depth, the quantitative similarity between predictions and observations is intriguing.

Finally, our work potentially opens ways to understand confirmation biases. Confirmation biases happen when people extensively sample too few alternatives, thus effectively seeking information for the same source. We have demonstrated that oversampling some alternatives and completely ignoring others is optimal in certain conditions. It remains to be seen, however, whether this is actually the optimal strategy under more general conditions or whether the oversampling strategy induces severely harmful biases in certain niches.

It is important to note that we have described the phenomenology of the breadth-depth dilemma in conditions where all alternatives are, *a priori*, equally good. Thus, ignoring a large fraction of options and the associated square root sampling law can only be the worst-case scenario, in the sense that if there are biases or knowledge that a subset of alternatives is initially better than the rest, then fewer number of alternatives should be sampled. This consideration reassures us in the conclusion that the number of alternatives that ought to be sampled is much smaller than sampling capacity, an observation that might turn to be of general validity in both decision-making setups as well as in terms of brain organization for cognition.

## Acknowledgements

This work is supported by the Howard Hughes Medical Institute (HHMI, ref 55008742), MINECO (Spain; BFU2017-85936-P) and ICREA Academia (2016) to R.M.-B.; by NIH (DA037229) to B.Y.H.; by a scholar award from the James S. McDonnell Foundation (grant# 220020462) to J.D.

## Data Availability

The data that support the findings of this study are publicly available at https://github.com/rmorenobote/breadth-depth-dilemma

## Code Availability

The codes used for analysis and to generate figures are available publicly at https://github.com/rmorenobote/breadth-depth-dilemma

## Bibliography

Averbeck, B.B. (2015). Theory of Choice in Bandit, Information Sampling and Foraging Tasks. PLOS Computational Biology 11, e1004164.

Awh, E., Barton, B., and Vogel, E.K. (2007). Visual Working Memory Represents a Fixed Number of Items Regardless of Complexity. Psychological Science 18, 622–628.

Bates, C.J., Lerch, R.A., Sims, C.R., and Jacobs, R.A. (2019). Adaptive allocation of human visual working memory capacity during statistical and categorical learning. Journal of Vision 19, 11.

Bechhofer, R.E., and Kulkarni, R.V. (1984). Closed sequential procedures for selecting the multinomial events which have the largest probabilities. Communications in Statistics-Theory and Methods 13, 2997–3031.

Bettman, J.R., Luce, M.F., and Payne, J.W. (1998). Constructive Consumer Choice Processes. Journal of Consumer Research 25, 187–217.

Brady, T.F., Konkle, T., and Alvarez, G.A. (2011). A review of visual memory capacity: Beyond individual items and toward structured representations. Journal of Vision 11, 4–4.

Brandstätter, E., Gigerenzer, G., and Hertwig, R. (2006). The priority heuristic: Making choices without trade-offs. Psychological Review 113, 409–432.

Busemeyer, J.R., Gluth, S., Rieskamp, J., and Turner, B.M. (2019). Cognitive and Neural Bases of Multi-Attribute, Multi-Alternative, Value-based Decisions. Trends in Cognitive Sciences 23, 251–263.

Chen, W., Hu, W., Li, F., Li, J., Liu, Y., and Lu, P. (2016). Combinatorial Multi-Armed Bandit with General Reward Functions. Advances in Neural Information Processing Systems 29 1659–1667.

Cisek, P., and Kalaska, J.F. (2010). Neural Mechanisms for Interacting with a World Full of Action Choices. Annual Review of Neuroscience 33, 269–298.

Cohen, J.D., McClure, S.M., and Yu, A.J. (2007). Should I stay or should I go? How the human brain manages the trade-off between exploitation and exploration. Philosophical Transactions of the Royal Society B: Biological Sciences 362, 933–942.

Costa, V.D., Mitz, A.R., and Averbeck, B.B. (2019). Subcortical Substrates of Explore-Exploit Decisions in Primates. Neuron 103, 533–545.e5.

Cowan, N., Elliott, E.M., Scott Saults, J., Morey, C.C., Mattox, S., Hismjatullina, A., and Conway, A.R.A. (2005). On the capacity of attention: Its estimation and its role in working memory and cognitive aptitudes. Cognitive Psychology 51, 42–100.

Daw, N.D., O’Doherty, J.P., Dayan, P., Seymour, B., and Dolan, R.J. (2006). Cortical substrates for exploratory decisions in humans. Nature 441, 876–879.

Desimone, R., and Duncan, J. (1995). Neural mechanisms of selective visual attention. Annu Rev Neurosci 18, 193–222.

Deutsch, J.A., and Deutsch, D. (1963). Attention: Some theoretical considerations. Psychological Review 70, 80–90.

Diehl, K., and Poynor, C. (2010). Great Expectations?! Assortment Size, Expectations, and Satisfaction. Journal of Marketing Research 47, 312–322.

Ebitz, R.B., Albarran, E., and Moore, T. (2018). Exploration Disrupts Choice-Predictive Signals and Alters Dynamics in Prefrontal Cortex. Neuron 97, 450–461.e9.

Gershman, S.J., Horvitz, E.J., and Tenenbaum, J.B. (2015). Computational rationality: A converging paradigm for intelligence in brains, minds, and machines. Science 349, 273–278.

Gigerenzer, G., and Gaissmaier, W. (2011). Heuristic Decision Making. Annual Review of Psychology 62, 451–482.

Gittins, J.C., Weber, R., and Glazebrook, K.D. (2011). Multi-armed bandit allocation indices (Hoboken, NJ: John Wiley & Sons).

Griffiths, T.L., Lieder, F., and Goodman, N.D. (2015). Rational Use of Cognitive Resources: Levels of Analysis Between the Computational and the Algorithmic. Topics in Cognitive Science 7, 217–229.

Gupta, S.S., and Liang, T. (1989). Selecting the best binomial population: parametric empirical Bayes approach. Journal of Statistical Planning and Inference 23, 21–31.

Halpert, H. (1958). Folklore: Breadth versus Depth. The Journal of American Folklore 71, 97.

Hauser, J.R., and Wernerfelt, B. (1990). An Evaluation Cost Model of Consideration Sets. Journal of Consumer Research 16, 393.

Hayden, B.Y., and Moreno-Bote, R. (2018). A neuronal theory of sequential economic choice. Brain and Neuroscience Advances 2, 239821281876667.

Hick, W.E. (1952). On the Rate of Gain of Information. Quarterly Journal of Experimental Psychology 4, 11–26.

Hills, T.T., Todd, P.M., and Goldstone, R.L. (2010). The central executive as a search process: Priming exploration and exploitation across domains. Journal of Experimental Psychology: General 139, 590–609.

Hills, T.T., Jones, M.N., and Todd, P.M. (2012). Optimal foraging in semantic memory. Psychological Review 119, 431–440.

Horowitz, E., and Sahni, S. (1978). Fundamentals of computer algorithms (Potomac, Md: Computer Science Press).

Iyengar, S.S., and Lepper, M.R. (2000). When choice is demotivating: Can one desire too much of a good thing? Journal of Personality and Social Psychology 79, 995–1006.

Koechlin, E., and Summerfield, C. (2007). An information theoretical approach to prefrontal executive function. Trends in Cognitive Sciences 11, 229–235.

Korf, R.E. (1985). Depth-first iterative-deepening. Artificial Intelligence 27, 97–109.

Krajbich, I., Armel, C., and Rangel, A. (2010). Visual fixations and the computation and comparison of value in simple choice. Nature Neuroscience 13, 1292–1298.

Lim, S.-L., O’Doherty, J.P., and Rangel, A. (2011). The Decision Value Computations in the vmPFC and Striatum Use a Relative Value Code That is Guided by Visual Attention. Journal of Neuroscience 31, 13214–13223.

Luck, S.J., and Vogel, E.K. (2013). Visual working memory capacity: from psychophysics and neurobiology to individual differences. Trends in Cognitive Sciences 17, 391–400.

Ma, W.J., Husain, M., and Bays, P.M. (2014). Changing concepts of working memory. Nature Neuroscience 17, 347–356.

Mehta, N., Rajiv, S., and Srinivasan, K. (2003). Price Uncertainty and Consumer Search: A Structural Model of Consideration Set Formation. Marketing Science 22, 58–84.

Miller, D.P. (1981). The Depth/Breadth Tradeoff in Hierarchical Computer Menus. Proceedings of the Human Factors Society Annual Meeting 25, 296–300.

Miller, G.A. (1956). The magical number seven, plus or minus two: some limits on our capacity for processing information. Psychological Review 63, 81–97.

Morgan, P., and Manning, R. (1985). Optimal Search. Econometrica 53, 923.

Norman, D.A., and Shallice, T. (1986). Attention to Action. In Consciousness and Self-Regulation, R.J. Davidson, G.E. Schwartz, and D. Shapiro, eds. (Boston, MA: Springer US), pp. 1–18.

Ratcliff, R., and Murdock, B.B. (1976). Retrieval processes in recognition memory. Psychological Review 83, 190–214.

Redish, A.D. (2016). Vicarious trial and error. Nature Reviews Neuroscience 17, 147–159.

Rich, E.L., and Wallis, J.D. (2016). Decoding subjective decisions from orbitofrontal cortex. Nat Neurosci 19, 973–980.

Roberts, J.H., and Lattin, J.M. (1991). Development and Testing of a Model of Consideration Set Composition. Journal of Marketing Research 28, 429–440.

Roe, R.M., Busemeyer, J.R., and Townsend, J.T. (2001). Multialternative decision field theory: A dynamic connectionst model of decision making. Psychological Review 108, 370–392.

Rushworth, M.F., Noonan, M.P., Boorman, E.D., Walton, M.E., and Behrens, T.E. (2011). Frontal cortex and reward-guided learning and decision-making. Neuron 70, 1054–1069.

Russell, S., and Wefald, E. (1991). Principles of metareasoning. Artificial Intelligence 49, 361–395.

Santos, B.D. los, Hortaçsu, A., and Wildenbeest, M.R. (2012). Testing Models of Consumer Search Using Data on Web Browsing and Purchasing Behavior. American Economic Review 102, 2955–2980.

Scheibehenne, B., Greifeneder, R., and Todd, P.M. (2010). Can There Ever Be Too Many Options? A Meta-Analytic Review of Choice Overload. Journal of Consumer Research 37, 409– 425.

Schwartz, M.S., Sadler, P.M., Sonnert, G., and Tai, R.H. (2009). Depth versus breadth: How content coverage in high school science courses relates to later success in college science coursework. Sci. Ed. 93, 798–826.

Shadlen, M.N., and Shohamy, D. (2016). Decision Making and Sequential Sampling from Memory. Neuron 90, 927–939.

Shenhav, A., Barrett, L.F., and Bar, M. (2013). Affective value and associative processing share a cortical substrate. Cognitive, Affective, & Behavioral Neuroscience 13, 46–59.

Shenhav, A., Musslick, S., Lieder, F., Kool, W., Griffiths, T.L., Cohen, J.D., and Botvinick, M.M. (2017). Toward a Rational and Mechanistic Account of Mental Effort. Annual Review of Neuroscience 40, 99–124.

Siegel, M., Buschman, T.J., and Miller, E.K. (2015). Cortical information flow during flexible sensorimotor decisions. Science 348, 1352–1355.

Simon, H.A. (1955). A Behavioral Model of Rational Choice. The Quarterly Journal of Economics 69, 99.

Simon, HA (1972). Theories of bounded rationality. Decision and Organization *Chapter 8*.

Sims, C.A. (2003). Implications of rational inattention. Journal of Monetary Economics 50, 665–690.

Sims, C.R. (2016). Rate–distortion theory and human perception. Cognition 152, 181–198.

Sleezer, B.J., Castagno, M.D., and Hayden, B.Y. (2016). Rule Encoding in Orbitofrontal Cortex and Striatum Guides Selection. J Neurosci 36, 11223–11237.

Sobel, M., and Huyett, M.J. (1957). Selecting the Best One of Several Binomial Populations. Bell System Technical Journal 36, 537–576.

Stigler, G.J. (1961). The Economics of Information. Journal of Political Economy 69, 213–225.

Tajima, S., Drugowitsch, J., and Pouget, A. (2016). Optimal policy for value-based decision-making. Nat Commun 7, 12400.

Timmermans, D. (1993). The impact of task complexity on information use in multi-attribute decision making. Journal of Behavioral Decision Making 6, 95–111.

Treisman, A.M. (1969). Strategies and models of selective attention. Psychological Review 76, 282–299.

Turner, S.F., Bettis, R.A., and Burton, R.M. (2002). Exploring Depth Versus Breadth in Knowledge Management Strategies. Computational & Mathematical Organization Theory 8, 49–73.

Tversky, A. (1972). Elimination by aspects: A theory of choice. Psychological Review 79, 281–299.

Usher, M., and McClelland, J.L. (2004). Loss Aversion and Inhibition in Dynamical Models of Multialternative Choice. Psychological Review 111, 757–769.

Vickery, T.J., Chun, M.M., and Lee, D. (2011). Ubiquity and Specificity of Reinforcement Signals throughout the Human Brain. Neuron 72, 166–177.

Wilson, R.C., Geana, A., White, J.M., Ludvig, E.A., and Cohen, J.D. (2014). Humans use directed and random exploration to solve the explore–exploit dilemma. Journal of Experimental Psychology: General 143, 2074–2081.

Yantis, S., and Johnston, J.C. (1990). On the locus of visual selection: Evidence from focused attention tasks. Journal of Experimental Psychology: Human Perception and Performance 16, 135–149.

Yoo, S.B.M., and Hayden, B.Y. (2018). Economic Choice as an Untangling of Options into Actions. Neuron 99, 434–447.

