## Supplementary material for "The breadth-depth dilemma in a finite capacity model of decision-making": Methods

#### 1 Finite capacity model

We consider a two-stage decision process in a multi-alternative decision-making problem modeled as a partially observable Markov decision process. There are $N$ alternatives, defined each by a Bernoulli random processes, whose trial by trial ($t = 1, ...$) outcomes follow $s_i^t \sim \mathrm{Bern}\,(p_i)$, $s_i^t \in \{0, 1\} = \{\mathrm{failure}, \mathrm{success}\}$, $i = 1, ..., N$. The outcomes are independently distributed for all trials $t$. The values of the success probabilities are unknown to the decision-maker, and follow a prior distribution $p_i \sim \mathrm{Beta}(\alpha, \beta)$ i.i.d. for all alternatives, with known hyperparameters $(\alpha, \beta)$. Allowed actions follow a two-stage decision process. In the first stage, the decision-maker can draw a total of $C = N$ samples at once, namely, a one-go decision is considered [1, 2, 3]. We consider the case where the total number of alternatives $N$ exhausts sampling capacity $C$, but the results are equivalent if the number of alternatives is larger than capacity, with the addition of more rejected or non-sampled alternatives. The action space is $\mathsf{A_L} = \{\vec{L} : L_i \geq 0\ \forall i, \sum_i L_i = C\}$, where $\vec{L} = (L_1, ..., L_N)$ is the number of samples drawn from each of the alternatives, with the constraint $\sum_i L_i = C$ (we often refer to the vector $\vec{L}$ as sample allocation). Note that the decision-maker can decide to sample the same alternative several times (i.e., $L_i > 1$ for some

$i$), and also decide not to sample from several alternatives (i.e., $L_i = 0$ for other $i$). In general, $M \leq C = N$ alternatives are sampled. If just a few alternatives are sampled ($M \sim 1$), many samples can be allocated to each. If $C$ alternatives are sampled, only one sample could be allocated to each of them. Outcomes of the samples from the sampled alternatives are revealed all at once, not sequentially. In the second stage of the decision-making process, after outcomes are observed, the decision-maker should decide what alternative to choose. We initially assume that it is only possible to choose only among the sampled alternatives. Thus, the action space in the second stage is defined by the set $\mathsf{A_C} = \{c : L_c > 0\}$ of size $M$, ordered as $\{c_1, ..., c_M\}$. The sufficient statistics of the outcomes of the Bernoulli processes to infer the success probabilities are the counts of successes for each of the $M$ sampled alternatives, $\vec{n} = (n_{c_1}, ..., n_{c_M})$, with $n_j = \sum_{t=1}^{L_{c_j}} s^t_{c_j}$, and thus the decision of what option to choose should be a function of those counts and on the sample allocation vector $\vec{L}$, which together constitute the information state of the decision process. The counts, conditioned on the success probabilities, follow $n_i \sim \text{Bin}(p_i, L_i)$. Note that the dimension of the vector $\vec{n}$ depends on the number of samples alternatives (those satisfying $L_i > 0$) and thus the consideration set changes sizes depending on the first stage decision.

We define the utility of a choice $i \in \mathsf{A_C}$ as the hidden value of the success probability of the corresponding Bernoulli variable, $U_i = p_i$. We assume that the decision-maker maximizes expected utility. This involves determining the optimal allocation of samples $\vec{L}^*$ to be used in the first stage followed by defining an optimal decision rule that selects one of the sampled alternatives based on $\vec{n}$. A decision rules maps an observation $\vec{n}$, given the allocation vector $\vec{L}$, into an element of the action space $\mathsf{A_C}$. By considering all possible decision rules, $\delta = \{\delta(\vec{n}, \vec{L}) : (\vec{n}, \vec{L}) \to \mathsf{A_C}\}$, we show in Sec. (5) that the optimal decision rule, $\delta^*(\vec{n}, \vec{L})$, is the one that selects always, for any sample allocation $\vec{L}$, the alternative with the maximum posterior mean success probability $\mathbb{E}(p_i|n_i, L_i) = \frac{n_i+\alpha}{L_i+\alpha+\beta}$, $i \in A_C$, or chooses any of the maximum ones if there are ties. Therefore, the expected utility for a given sample allocation $\vec{L}$ following the optimal decision rule is

$$U(\vec{L}) = \sum_{\vec{n}} p\left(\vec{n}|\vec{L}, \alpha, \beta\right) \max_{i \in \mathsf{A_C}} \left(\frac{n_i + \alpha}{L_i + \alpha + \beta}\right), \tag{1}$$

where the joint posterior over $\vec{n}$ factorizes into beta-binomial distributions as $p\left(\vec{n}|\vec{L}, \alpha, \beta\right) = \prod_{i \in \mathsf{A_C}} \mathrm{Bb}(n_i|L_i, \alpha, \beta)$. Then, the optimal sample allocation $\vec{L}^*$ equals

$$\vec{L}^* = \arg\max_{\vec{L} \in \mathsf{A_L}} U(\vec{L}) = \max_{\vec{L} \in \mathsf{A_L}} \sum_{\vec{n}} p\left(\vec{n}|\vec{L}, \alpha, \beta\right) \max_{i \in \mathsf{A_C}} \left(\frac{n_i + \alpha}{L_i + \alpha + \beta}\right), \tag{2}$$

and the corresponding maximum expected utility becomes

$$U^* = \max_{\vec{L} \in \mathsf{A_L}} U(\vec{L}). \tag{3}$$

Finding the optimal solution in Eq. (2) is hard because of the large number of sample allocations that it is possible to form out of $C$ samples. The number of unique partitions of $C$ samples equals the number of integer partitions of $C$ (not to be confused with the Bell number), for which we are not aware of simple exact expressions. We should only consider unique partitions because all the alternatives are initially (before sampling) indistinguishable. Therefore, without loss of generality, we can always assume that we sample the alternatives by using the sample allocation $\vec{L} \in \mathsf{A_L}$ where we impose the additional constraint that $L_i \geq L_{i+1}$ for $i = 1, ..., N-1$. That is, we sample the first alternative with more or the same number of samples as the second alternative, the second alternative with more or the same number of samples as the third one, and so forth. We describe a gradient descent approach bellow in Sec. (4) to find the optimal sample allocation exactly for small capacity $C$ and approximately for large capacity. To find useful analytical expressions for Eqs. (2, 3), we restrict ourselves further by first looking for optimal even sample allocations, that is, allocation of samples across $M \leq C$ options with the same number of samples $L$ per alternative. Optimal even sample allocation across alternatives is discussed in Sec. (2).

### 2 Analytical expressions for optimal even sample allocation

Because the space of actions $\mathsf{A_L} = \{\vec{L} : L_i \geq 0 \ \forall i, \sum_i L_i = C\}$ is very large, we restrict ourselves to a subset of possible actions, consisting in dividing the capacity $C$ into $M$ alternatives equally sampled with $L$ samples each. Without loss of generality, we assume that we sample the first $M$ alternatives and we ignore the rest of $N - M$ alternatives. Even splitting of the capacity is only possible if $C = M \times L$ holds exactly, so we will only examine the pairs $(M, L)$ that satisfy that condition. The advantage of working in this subset of actions is that it is possible to obtain useful, exact analytical expression that will reveal non-trivial properties of the decision process. Methods for finding globally optimal sample allocation strategies are provided in Sec. (4). In the main results we also show that optimal sample allocations are not greatly better than the optimal even ones, so that even sample allocation is close-to-optimal. For an even capacity split, the optimal $L^*$ under the constraint $C = ML$ can be obtained by specializing Eq. (2) to this case as

$$L^* = \arg\max_L \sum_{\vec{n}} \prod_{j=1}^{M} p\left(n_j|L,\alpha,\beta\right) \max_i \left(\frac{n_i+\alpha}{L+\alpha+\beta}\right), \tag{4}$$

where $i \in \{1,...,M\}$ and $p\left(n_j|L,\alpha,\beta\right) = \mathrm{Bb}(n_j|L,\alpha,\beta)$. Naturally, the optimal number of alternatives to be sampled is $M^* = C/L^*$

A particularly simple expression results from the case $\alpha = \beta = 1$, corresponding to a uniform prior over the success probabilities of the Bernoulli variables. This is because $p\left(n_j|L,1,1\right) = \mathrm{Bb}(n_j|L,1,1) = \frac{1}{L+1}$, thus becoming a discrete uniform distribution over $n_j \in \{0,...,L\}$, independent of $n_j$. Then, replacing this expression in Eq. (4), the optimal even sample allocation simplifies to

$$\begin{aligned}
L^* &= \arg\max_L U(L), \\
U(L) &= \frac{1}{(L+1)^M} \sum_{n_1,...,n_M=0}^{L} \max_i \left(\frac{n_i+1}{L+2}\right) && (5)\\
&= \frac{1}{(L+1)^M(L+2)} \left((L+1)^M + \sum_{n_1,...,n_M=0}^{L} \max(n_1,...,n_M)\right) \\
&= \frac{1}{(L+1)^M(L+2)} \left((L+1)^M + \sum_{s=0}^{L} \left((s+1)^M - s^M\right) s\right) && (6)\\
&= 1 - \frac{\sum_{s=0}^{L}(s+1)^M}{(L+1)^M(L+2)}, && (7)
\end{aligned}$$

with $M = C/L$. Eq. (6) in the derivation results from realizing that the sum over $\max_i(n_i)$ contains exactly $1^M - 0$ zeros, $2^M - 1$ ones, $3^M - 2^M$ twos, etc. The sum in Eq. (7) is the sum of the $M-th$ powers of the first $L+1$ integers, and it can be computed using Faulhaber's formula. Eq. (7) confirms the intuition that the expected utility $U(L)$ for any $L$ is smaller than one. Finally, the optimal number of evenly allocated samples (over the sampled options) can be written as

$$L^* = \arg\min_L \frac{\sum_{s=0}^{L}(s+1)^M}{(L+1)^M(L+2)} \tag{8}$$

It is interesting to examine some limits of Eq. (7) by relaxing the constraint $C = M \times L$. For large $M$ and $L = 1$, the expected utility in Eq. (7) becomes $\lim_{M\to\infty} U \to \frac{2}{3}$. This observation is not surprising, as when a very large number of alternatives is sampled with just one sample, it is very likely that at least one of them will have a successful outcome. Therefore, the expected utility of that alternative under the uniform prior will be $\frac{2}{3}$. This limit is visible in the rightmost point of Fig. 2a. In the opposite scenario, when only one alternative is sampled, $M = 1$, then the expected utility is $\frac{1}{2}$ for all $L$. That is, if just one alternative is sampled, then the expected probability of success of the sampled alternative is $\frac{1}{2}$, which equals the prior mean. This limit is visible in the leftmost point of Fig. 2a.

A more general way of performing the integrals involved in Eq. (4) is by using cumulative distribution function of the beta-binomial distributions, $\phi(n|L,\alpha,\beta) = \sum_{m\leq n} \mathrm{Bb}(m|L,\alpha,\beta)$, and write the optimal even sample allocation in Eq. (4) in terms of extreme value distribution $\phi^M(n) - \phi^M(n-1)$ as

$$L^* = \arg\max_L \sum_{n=0}^{L} \left[\phi^M\left(n|L,\alpha,\beta\right) - \phi^M\left(n-1|L,\alpha,\beta\right)\right] \left(\frac{n+\alpha}{L+\alpha+\beta}\right), \tag{9}$$

Note that the extreme value distribution $\phi^M(n_{max}) - \phi^M(n_{max} - 1)$ is the distribution of $n_{max} = \max(n_1, ..., n_M)$ where $\vec{n}$ follows the above factorized beta-binomial distribution. In other words, the extreme value distribution for $n_{max}$ is the probability that no alternative has more than $n_{max}$ successful samples (hence the first term $\phi^M(n_{max})$) but removing the cases where there is no alternative with more than $n_{max}-1$ successful samples (hence the second negative term $\phi^M(n_{max}-1)$). For the uniform prior case, $\alpha = \beta = 1$, we recover Eq. (8), for which the cumulative can be exactly computed. For arbitrary values of $\alpha$ and $\beta$, Eq. (9) is solved numerically. These solutions are used in Fig. 2d.

The general Eq. (2) valid for any allocation of samples, and the specific Eq. (9) valid for even sample allocations, assume that a choice is made from the sampled alternatives, while non-sampled alternatives are excluded from the choice set. However, if none of the sampled alternatives turns to be good ones (e.g., because $n_i \ll L_i$ for $i \in \mathsf{A_C}$), then it would be better to choose randomly from any of the non-sampled alternatives. This is particularly so if the expected utility of any of the sampled alternatives, $\frac{n_i+\alpha}{L_i+\alpha+\beta}$, is smaller than $\frac{\alpha}{\alpha+\beta}$, which is the default expected utility of the non-sampled alternatives given that the success probabilities are drawn from a $\mathrm{B}(\alpha, \beta)$. It is straighforward to generalize these results by adding a default alternative, assumed to have utility $p_0$. In this case, the optimal even allocation of samples obeys

$$L^* = \arg\max_L \sum_{n=0}^{L} \left(\phi^M\left(n|L,\alpha,\beta\right) - \phi^M\left(n-1|L,\alpha,\beta\right)\right) \max\left(\frac{n+\alpha}{L+\alpha+\beta},\ p_0\right). \tag{10}$$

### 3 Asymptotic behavior for large capacity: the square root sampling law

It is possible to derive an approximation for the limiting behavior of the optimal number of sampled alternatives $M^*$ and their associated optimal number of samples per alternative $L^*$ by using Eq. (5) for large capacity $C$ in the case of the uniform prior distribution. For large capacity $C$, we assume that $L^*$ grows to infinity. This assumption is confirmed later, when the asymptotic optimal $L^*$ is derived. If $L$ is large, then Eq. (5) can be approximated by

$$\begin{aligned} U(L) &= \frac{1}{(L+1)^M} \sum_{n_1,...,n_M=0}^{L} \max_i \left( \frac{n_i+1}{L+2} \right) \\ &= \frac{1}{(L+2)} \left( 1 + \frac{1}{(L+1)^M} \sum_{n_1,...,n_M=0}^{L} \max(n_1,...,n_M) \right) \\ &\approx \frac{1}{(L+2)} \left( 1 + L \int_0^1 dx_1 ... \int_0^1 dx_M \max(x_1,...,x_M) \right), \end{aligned} \qquad (11)$$

where the sum in the second equation has been approximated in the third equation by an integral in the interval $[0,1]^M$ over a uniform distribution by using the transformation $n_i = Lx_i$ for $i = 1, ..., M$. The continuous approximation is valid when $L$ is large, as assumed, since then the transformation delivers values of $x_i$ that are dense in the unit interval. The integral can be rewritten as

$$\int_0^1 dx_1 ... \int_0^1 dx_M \max(x_1,...,x_M) = \int_0^1 dx_{max} x_{max} f(x_{max}),$$

where we have defined the extreme value $x_{max} = \max(x_1, ..., x_M)$. The extreme value follows the extreme value distribution $f(x_{max}) = (F(x_{max})^M)' =$

$Mx_{max}^{M-1}$, where we have used that $F(x) = x$ is the cumulative of the continuous uniform distribution in $[0,1]$. Therefore,

$$
\begin{aligned}
U(L) &\approx \frac{1}{(L+2)}\left(1 + L\int_0^1 dx_{max} M x_{max}^M\right) \\
&= \frac{1}{(L+2)}\left(1 + \frac{ML}{M+1}\right). \qquad (12)
\end{aligned}
$$

Finally, by maximizing $U(L)$ as a function of $L$ with the constraint $C = ML$ we obtain the asymptotic optimal number of sampled alternatives $M^*$ and optimal number of samples per sampled alternative $L^*$

$$
\lim_{C\to\infty} M^* = \sqrt{C}, \quad \lim_{C\to\infty} L^* = \sqrt{C},
$$

which corresponds to the square root sampling law.

In the above derivation we have assumed that $L^*$ grows with $C$. To see that this corresponds to the only valid assumption to obtain $L^*$, let us assume now that $L^*$ does not grow with $C$, that is, it is a constant or decreases with $C$. For any fixed value $L$, using Eq. (7) we see that $U(L) \leq 1 - 1/(L+2)$. This utility is smaller than the one obtained by using the square root law, which converges to 1, as can be easily derived from Eq. (12). Therefore, the square root law delivers the highest utility.

### 4 Optimal sample allocation

For low capacity $C \leq 7$ we found the globally optimal sample allocation strategy by exhaustive search over all possible sample allocations. For larger capacity, we searched the optimal sample allocation by using Monte Carlo gradient descent. With this method, we confirmed that for values up to $C \leq 20$ the globally optimal sample allocations were correct up to a precision in expected utility of $10^{-4}$.

We started the algorithm by selecting the optimal even sample allocation if $C \leq 7$, and the square root law if the capacity was larger (we considered the possibility that the resulting square root was not an integer, and thus we allocated the residual number of samples to a randomly chosen additional alternative; we call this allocation scheme 'even allocation'). At every iteration, we computed the expected utility of the current best sample allocation $\vec{L}$ through a Monte Carlo simulation of the Bernoulli variables and averaging utility over $4 \times 10^5$ repetitions for $C \leq 20$ and $5 \times 10^4$ for larger capacity values. A perturbed sample allocation was proposed by randomly selecting two alternatives. One sample was removed from the first alternative and added to the second one, but only if the first alternative had already assigned at least one sample. To exploit symmetry, we only consider changes of one sample from one alternative $i$ to another $j > i$ if $L_{j-1} \geq L_j$ and $L_i \geq L_j$. If $j < i$, there were not restrictions.

With the proposed sample allocation, we computed the expected utility using the same Monte Carlo method. If the new expected value was larger than the previous one, then the proposed perturbed sample allocation became the current best sample allocation. This process was iterated $2 \times 10^4$ times for $C \leq 20$ and $3 \times 10^3$ for larger capacity values. Because at each iteration we reevaluate the expected value of the current best sample allocation, we avoid the possibility of getting stuck in a random fluctuation leading to a spuriously large expected value. The Monte Carlo gradient descent method found optimal sample allocations that were identical to those found with the exhaustive search for low capacity. We confirmed that the optimal sample allocations found were stable against different random number seeds and initial conditions. Figs. 3 and 4 use the above method. Percentage points increase in Fig. 4b is computed as $100(U^* - U_{even})/U_{even}$, where $U^*$ is the utility estimate of the globally optimal allocation and $U_{even}$ is the estimate of the even allocation.

We also used another version of the Monte Carlo gradient descent that avoided sampling the Bernoulli variables to estimate expected utility. This method was used to confirm robustness of the previous results. In this Markov Chain Monte Carlo estimation of utility, we define

$$U^* = \max_{\vec{L}} \sum_{\vec{n}} p(\vec{n}|\vec{L}, \alpha, \beta) \max_i \left( \frac{n_i + \alpha}{L_i + \alpha + \beta} \right). \qquad (13)$$

We thus can design a Markov Chain Monte Carlo method to sample from the probability distribution

$$p(\vec{n}|\vec{L}, \alpha, \beta) = \prod_j Bb(n_j|\vec{L}, \alpha, \beta)$$

appearing in the sum of Eq. (13) as follows (these samples can be then used to approximate the sum). Detailed balance imposes that the probability of transitioning from a state with $\vec{n}$ to $\vec{n}'$ is the same as the converse,

$$P_{\vec{n},\vec{n}'}\, p(\vec{n}|\vec{L}, \alpha, \beta) = P_{\vec{n}',\vec{n}}\, p(\vec{n}'|\vec{L}, \alpha, \beta).$$

By proposing a change to a single arm $n'_j = n_j \pm 1$, we can get a simple expression for the acceptance rate $r(\vec{n} \to \vec{n}')$. If $n'_j = n_j + 1$ the acceptance rate is

$$r(\vec{n} \to \vec{n}') = \min \left( 1, \frac{(n_j + \alpha)(L_j - n_j)}{(L_j - n_j + \beta + 1)(n_j + 1)} \right),$$

while if $n'_j = n_j - 1$, it becomes

$$r(\vec{n} \to \vec{n}') = \min\left(1, \frac{(L_j - n_j + \beta)n_j}{(n_j + \alpha - 1)(L_j - n_j)}\right),$$

where we have made use of the Metropolis-Hastings algorithm. These two changes are proposed with equal probability and randomly across all the options. Utilities are estimated using $10^6$ samples. The search over $\vec{L}$ is made using $50 \times C$ iterations for $C \leq 50$ and 2500 iterations for $50 < C < 5000$.

### 5 Consistency

Perhaps intuitively, but wrongly, we might assume that by always opting for the alternative with larger number of successful outcomes (larger $n_i$ in Eq. (2)), this would result in 'cherry picking', that is, in selected a spuriously good option. This, in turn, would mean that we would obtain a reward that is lower than the expected utility in Eq. (3). Here we show, however, that the decision rule of choosing always the alternative with the highest posterior mean is both optimal and delivers on average a reward that is equal to the expected utility. This is a well-known result in statistical decision theory [4, 5, 6]. Here we show the derivation for completeness.

Consider any possible decision rule $\vec{d} = \delta(\vec{n})$ that assigns the counts of successes for the $M$ sampled alternatives, $\vec{n}$, to a decision $\vec{d} \equiv \vec{d}(\vec{n}) = (d_{c_1}(\vec{n}), ..., d_{c_M}(\vec{n}))$, encoded as a one-hot vector of length $M$ (i.e., $d_{c_i} = 1$ if alternative $c_i$ is chosen, and $d_{c_i} = 0$ otherwise; we omit the potential dependence of the decision rule on $\vec{L}$ to avoid cluttered notation). If the success probabilities of the sampled alternatives, $\vec{p}$, are known, then by using the decision rule $\delta$ the decision-maker would have an expected utility

$$U(\vec{p}, \vec{L}, \delta) = \sum_{\vec{n}} \prod_{i \in \mathsf{A_C}} \mathrm{Bin}(n_i | L_i, p_i)\, p_i^{d_i(\vec{n})},$$

where $\vec{L}$ is the allocated number of samples over the alternatives. Note that the expected utility is an average over the values of the chosen $p_i$ given the decision rule averaged across all possible outcomes given the allocated number of samples over alternatives. As probabilities are unknown, they are marginalized out with their prior beta distributions, resulting in the overall expected utility

$$U(\vec{L}, \delta) = \sum_{\vec{n}} \prod_{i \in \mathsf{A_C}} \frac{\Gamma(L_i + 1)}{\Gamma(n_i + 1)\Gamma(L_i - n_i + 1)} \frac{\Gamma(\alpha + \beta)}{\Gamma(\alpha)\Gamma(\beta)} \times \frac{\Gamma(n_i + \alpha + d_i)\Gamma(L_i - n_i + \beta)}{\Gamma(L_i + \alpha + \beta + d_i)}. \tag{14}$$

We note that for each term in the sum over $\vec{n}$, there is only one value of $i$ for which $d_i = 1$ in the product, while $d_j = 0$ for $j \neq i$. The term $i$ in the product with $d_i = 1$ gives an extra factor $\frac{n_i+\alpha}{L_i+\alpha+\beta}$ (by expanding the gamma functions just one step) that is not present in the product terms with $d_j = 0$. Therefore, the product is maximized iff $d_i = 1$ for the alternative $i$ with maximum $\frac{n_i+\alpha}{L_i+\alpha+\beta}$ (if the maximum is not unique, any alternative with the maximum value will give exactly the same result). This result proves that the optimal decision rule $\delta^*$ is the one that chooses always the alternative with the highest posterior expected utility given $\vec{n}$.

Now, we can show that for the optimal decision rule $\delta^*$, the expected utility is the same as that in Eq. (3). We can rewrite Eq. (14) as

$$U(\vec{L}, \delta^*) = \sum_{\vec{n}} \max_{i \in \mathsf{A_C}} \left( \frac{n_i + \alpha}{L + \alpha + \beta} \right) \prod_{i \in \mathsf{A_C}} \frac{\Gamma(L_i + 1)}{\Gamma(n_i + 1)\Gamma(L_i - n_i + 1)} \frac{\Gamma(\alpha + \beta)}{\Gamma(\alpha)\Gamma(\beta)} \times \frac{\Gamma(n_i + \alpha)\Gamma(L_i - n_i + \beta)}{\Gamma(L_i + \alpha + \beta)},$$

which is identical to the maximum expected utility $U(\vec{L})$ in Eq. (3), that is, $U(\vec{L}, \delta^*) = U(\vec{L})$. This shows that 'cherry picking' is optimal.

### References


[1] Bechhofer, R. E., Kulkarni, R. V. Closed sequential procedures for selecting the multinomial events which have the largest probabilities. Communications in Statistics - Theory and Methods 13, 2997–3031 (1984).

[2] Gupta, S. S., Liang, T. Selecting the best binomial population: parametric empirical Bayes approach. Journal of Statistical Planning and Inference 23, 21–31 (1989).

[3] Sobel, M., Huyett, M. J. Selecting the Best One of Several Binomial Populations. Bell System Technical Journal 36, 537–576 (1957).

[4] Moreno-Bote, R. Decision Confidence and Uncertainty in Diffusion Models with Partially Correlated Neuronal Integrators. Neural Computation 22, 1786–1811 (2010).

[5] Beck, J. M. et al. Probabilistic population codes for Bayesian decision making. Neuron 60, 1142–52 (2008).

[6] Berger, J. O. Statistical Decision Theory and Bayesian Analysis. (Springer New York, 1985).
